# Identification of a filamentous form of kunitz protease inhibitor in *Asteraceae*

**DOI:** 10.1101/2021.01.22.427805

**Authors:** Sara Bratsch, Neil Olszewski, Benham Lockhart

## Abstract

Filamentous structures were observed in purified extracts from chrysanthemum, gerbera, sunflower and zinnia. When purified filament proteins were subjected to SDS-PAGE, the major protein associated with filaments from all three species has an apparent molecular mass of ≈25 kDa. Protein bands from chrysanthemum, gerbera, and zinnia were subjected to N-terminal protein sequencing while proteins from sunflower were sequenced by CID MS/MS. All of the sequences shared highest similarity to the kunitz trypsin inhibitor family. The sequencing results indicated that the proteins lacked the signal sequences. We tested the gerbera filament protein for glycosylation and found that it was a glycoprotein. Together these results indicate that the filaments are composed of mature KTI protein. This is the first report of a KTI assembling into filaments and the first report of a filament forming *Asteraceae* enzyme.

## Introduction

The *Asteraceae* family is one of the largest families of flowering plants belonging to the *Angiospermae* group (Jeffrey 2007). The *Asteraceae* group name comes from the type genus *Aster*, Greek for star, and is referring to the star-like flowers of the family. The *Asteraceae* family is natively found throughout the world in all habitats except Antarctica and the extreme Arctic. Economically important members include the sunflower, *Helianthus annuus*, used for trans-fat free oil and a whole seed crop; pyrethrum, *Chrysanthemum cineariifolium* and *C. coccineum*, from which the natural insecticide pyrethrin is extracted (Casida 1980); stevia, *Stevia rebaudiana*, a low calorie sweetener (Brandle et al 1998); lettuce, *Lactuca sativa*, a leafy vegetable; Jerusalem artichoke or sunchoke, *H. tuberosus*, a tuberous root vegetable; and others. Numerous genera are used horticulturally for cut flowers and ornamental plants like the chrysanthemum, *Chrysanthemum* spp.; coneflower, *Echinaceae* spp.; dahlia, *Dahlia* spp.; gerbera daisy, *Gerbera* spp.; pot marigold, *Calendula offinalis*; sunflower, *Helianthus* spp.; and zinnia, *Zinnia* spp.

Here we report the discovery of proteinaceous filaments in several members of the *Asteraceae*. In addition to cytoskeleton enzymes such as actin and tubulin, over 100 enzymes that self-assemble into filaments have been identified in both the Prokaryota and the Eukaryota (Park and Horton, 2019). These filaments can have different structures and in some cases a single enzyme can form filaments with different structures. While much remains to be learned about these filaments it is known that assembly into a filament can alter enzymatic activity. Enzyme filaments were first observed in plants by electron microscopy in 1965 (Gunning, 1965). These filaments are composed of β-glucosidase. In plants three enzymes including β-glucosidase, ribonucleotide reductase, and IRE1 have been identified forming filamentous structures (Park & Horton 2019).

Kunitz soybean trypsin inhibitor (KTI) proteins are widely distributed throughout plants and are often encoded by large gene families ranging in size from two to more than 50 genes (Fischer et al 2015; Islam et al 2015). KTIs are plant storage proteins with various functions that can be found in both seeds and vegetative tissues (de Souza Cândido et al. 2011). These proteins are produced at the cytoplasmic surface of the rough endoplasmic reticulum (de Souza Cândido et al. 2011). Most KTIs have an N-terminal signal sequence which directs the proteins into the ER and ultimately deposited in storage vacuoles. During this process the signal peptide is removed and the protein is glycosylated (Muntz 1998, Ee et al. 2011). KTIs collectively inhibit a wide range of proteolytic enzymes (Fischer et al 2015; Heibges et al 2003; Major & Constabel 2008). In addition, some KTIs have invertase inhibitory activity (Glaczinski et al 2002). Expression of KTIs can be constitutive or induced by herbivory, wounding, pathogens, abiotic stresses and plant hormones (Kang et al 2002; Kim et al 2003; Li et al 2008). KTIs can protect plants from herbivory, pathogens, and some KTIs have a role in plant development, since reducing their expression affects leaf shape, shoot growth and root growth (Islam et al 2015; Li et al 2008).

Here we report the identification and characterization of novel filamentous structures in members of the family *Asteraceae* including: chrysanthemum (*Chrysanthemum* sp.), coneflower (*Echinaceae* spp.), gerbera (*Gerbera jamesonii, G. hybrida*), sunflower (*Helianthus anuus*), and zinnia (*Zinnia hybrida*). Two different protein sequencing methods revealed that these filamentous structures were composed of a protein belonging to the soybean kunitz trypsin inhibitor family.

## Materials and methods

### Filament Purification

Filaments were purified for characterization by transmission electron microscopy from chrysanthemum (*Chrysanthemum* spp.), coneflower (*Echinacea* spp.), gerbera (*Gerbera jamesonii, G. hybrida*), sunflower (*Helianthus annuus*), or zinnia (*Zinnia hybrida*) leaf tissue using a differential centrifugation-based method. Filaments were extracted from leaf tissue that was frozen in liquid nitrogen, ground with a mortar and pestle, and mixed with 15 mL of extraction buffer (500 mM NaPO_4_, 1 M urea, 5% (w/v) polyvinylpyrrolidone, 0.02% (w/v) NaN_3_, 0.5% (v/v) 2-mercaptoethanol pH 7.5). This mixture was filtered using polyester fabric and centrifuged at 19,800 g*_max_* for 15 min at 10°C. The supernatant was removed and mixed with one mL of 25% (v/v) Triton X-100 by vortexing which was layered over five mL of 30% (w/v) sucrose in 100 mM NaPO_4_ pH 7.5 and subsequently centrifuged at 109,000 g_max_ for 2 hours at 10°C. The supernatant was discarded. The pellet was suspended with 100 μL of 100 mM NaPO_4_ pH 7.5. The solution was emulsified by vortexing with 100 μL chloroform and 20 μL 25% Triton X-100 and was centrifuged for 10 minutes at 26,400 g_*max*_ at room temperature (RT). The upper aqueous phase was used for TEM examination.

Filament preparations used to determine the identity of the filament proteins were further purified by cesium chloride isopycnic density-gradient centrifugation. The aqueous upper phase from above was applied to a preformed 0-30% cesium chloride gradient with 10% (w/w) sucrose in 100 mM NaPO_4_ pH 7.0. The gradient was centrifuged for 4.5 hours at 116,000 g*_max_* at 10°C. The gradient was fractioned into 500 μL fractions. Each fraction was examined by TEM as described below and fractions containing highly enriched filaments were pooled. The pooled fractions were diluted four-fold with 100 mM NaPO_4_ pH 7.5, and concentrated by centrifuging at 136,000 g*_max_* for 2 hours. The pellet was resuspended in 100 mM NaPO_4_, pH 7.5, containing 0.5% (v/v) 2-mercaptoethanol and examined by TEM as described below.

### Electron Microscopy

Two μL of the sample was mounted onto 0.25% Formvar and carbon coated 200 mesh copper grids (Electron Microscopy Sciences, USA). The grids were examined using a Phillips CM12 transmission electron microscope (University Imaging Centers, University of Minnesota) after negative staining with 2% (w/v) sodium phosphotungstate (PTA), pH 7.0, containing bacitracin at 100 μg/ml. The average diameter of filaments was calculated by measuring 50 particles each from coneflower, gerbera, pyrethrum, sunflower, and zinnia using Fiji (Schindelin et al. 2012).

### SDS-PAGE

Purified filaments were resuspended in SDS-PAGE sample buffer and filament proteins were subjected to SDS-PAGE on 4-20% TruPAGE Precast Gels (Sigma-Aldrich, USA). Gels were stained with Coomassie brilliant blue G-250 stain (Sigma-Aldrich, USA).

### Staining of glycoproteins

Filaments extracted from gerbera were subjected to SDS-PAGE and then the Pierce^™^ Glycoprotein Staining Kit (ThermoFisher Scientific, USA) was used as directed. To visualize nonglycosylated proteins the gel was also stained with Coomassie brilliant blue.

### N-terminal protein sequencing

N-terminal protein sequencing was performed by the Protein Facility of the Iowa State University Office of Biotechnology. Purified filament samples were submitted for analysis. For analysis the proteins were separated on SDS-PAGE and transferred to polyvinylidene difluoride membrane (PVDF). The membrane was stained with Coomassie brilliant blue R-250: 40% methanol: 10% acetic acid for 5 minutes and destained. Detectable protein bands were excised for sequencing. Samples were sequenced by Edman Degradation using a Perkin Elmer 494 Procise Protein/Peptide Sequencer with an on-line 140C PTH Amino Acid Analyzer (Applied Biosystems, Inc).

### Cloning and sequencing the sunflower kunitz protease inhibitor gene

The N-terminal sequence from chrysanthemum was used for a tBlastn search of the NCBI EST database. This search identified chrysanthemum EST sequence DK939406.1 but the sequence did not cover the entire coding region. The EST was used as a starting point to assemble a putative sequence of the coding region from other sequences in the NCBI database. Two primers, sunflower_KTI_F (5’- ATGAAGATCACATTGTCTTTCATTTTCTTA −3’) and sunflower_KTI _R (5’ – CTACTCAGATTGAACAGAAGCCAC- 3’were designed and used in a PCR reaction with total DNA extracted (DNeasy Plant Mini Kit, Qiagen, Valencia CA USA) from sunflower leaf tissue. The PCR reaction was performed using Platinum SuperFi DNA Polymerase I (ThermoFisher Scientific, USA) and a cycling program of 95 °C for 5 minutes followed by 25 cycles of 95 °C for 30 seconds, 60 °C for 30 seconds, 72 °C for 30 seconds, with a final extension at 95 °C for 10 minutes. The PCR product was purified (Pure link Quick PCR purification kit, Invitrogen USA), cloned and both strands were sequenced (UMGC, St. Paul USA).

### CID MS/MS sequencing

Proteins associated with purified sunflower filaments were submitted to the Center for Mass Spectrometry & Proteomics (CMSP) Facility (University of Minnesota) for peptide sequencing by mass spectrometry. Briefly, the purified filaments were denatured and analyzed by SDS-PAGE using Criterion^™^ Precast Gels (Bio-Rad). The gel was stained with Imperial Protein Stain (ThermoFisher Scientific). Protein bands of interest were excised to prepare for sequencing and destained (Drbal et al 2001). Proteolytic digestion was completed on the excised protein bands by in-gel trypsin digest using a protocol adapted from Shevchenko et al (1996). The samples were then desalted by STop And Go Extraction (STAGE) TIPS desalting procedure (Rappsilber & Mann, 2003). Mass spectrometry (MS) was completed as described in Lin-Moshier et al (2013).

Peaks Studio 6.0 build 20120620 (Bioinformatics Solutions Inc.) software package was used for interpretation of tandem MS and protein inference (Ma et al 2003). Search parameters included UniProt database (Campanula genera taxonomy ID 91882, accessed Jan 6 2016) concatenated with the common lab contaminants (cRAP) database (http://www.thegpm.org/crap/) and the translated KTI coding region obtained by PCR. In addition, an artificial proteome was created by taking the sunflower transcriptome assembled by Trinity (https://www.sunflowergenome.org/transcriptome.html) and translating it. This artificial proteome was also used in the search parameters. The remaining search parameters for Peaks were followed as described in Dahlin et al (2015).

## Results

Filaments extracted from chrysanthemum (*Chrysanthemum* spp.), coneflower (*Echinacea* spp.), gerbera (*Gerbera jamesonii, G. hybrida*), sunflower (*Helianthus annuus*), and zinnia (*Zinnia hybrida*) leaf tissue are shown in Figure 1. The filaments had mean diameters ranging from 7.6 to 8.9 nm (Table 1) with a mean across all species of 8.5 nm and variable length ranging up to several thousand nm.

**Figure 1.**
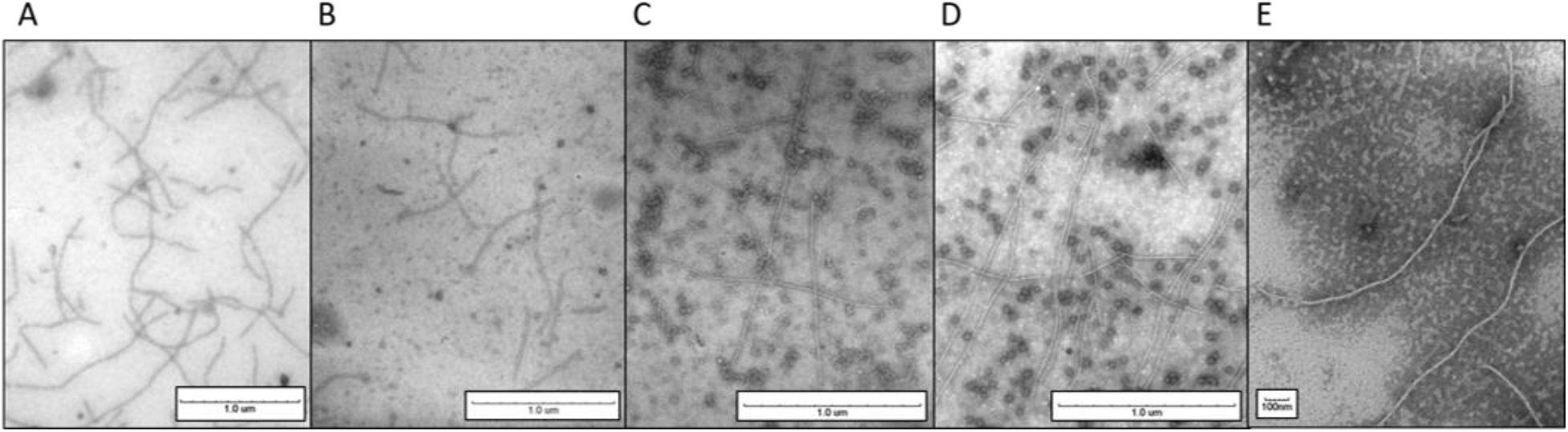
Transmission electron micrographs of filaments from A, sunflower (25,000X); B, coneflower (31,000X); C, chrysanthemum (40,000x); D, gerbera daisy (40,000X); and E, zinnia (53,000x). Filaments in B, C and D were partially purified and the filaments in A and E were further purified by centrifugation on a CsCl-sucrose gradient. The filaments were stained with 2% phosphotungstic acid.

**Table 1.** Filament diameter obtained by measuring 50 filaments each from coneflower, gerbera, chrysanthemum, sunflower, and zinnia.

|  | chrysanthemum | coneflower | gerbera | sunflower | zinnia |
| --- | --- | --- | --- | --- | --- |
| **Mean** | 8.57 | 8.94 | 7.64 | 8.91 | 8.39 |
| **SD** | 1.16 | 0.98 | 0.92 | 0.88 | 1.01 |
| **Min** | 6.38 | 7.05 | 6.10 | 7.41 | 6.72 |
| **Max** | 10.47 | 10.63 | 9.69 | 10.70 | 10.41 |

Proteins from sunflower, gerbera, and zinnia filament preparations were resolved by SDS-PAGE (Figure 2). All of the filament preparations had a major protein that migrated with a molecular mass between 20 and 25kDa. Each preparation contained proteins with other molecular masses that were variably present in the different preparations. For example, the ≈10kDa and ≈60 kDa proteins present in the preparation shown in Figure 2B are far less abundant or absent from the preparations shown in Figure 2A.

**Figure 2.**
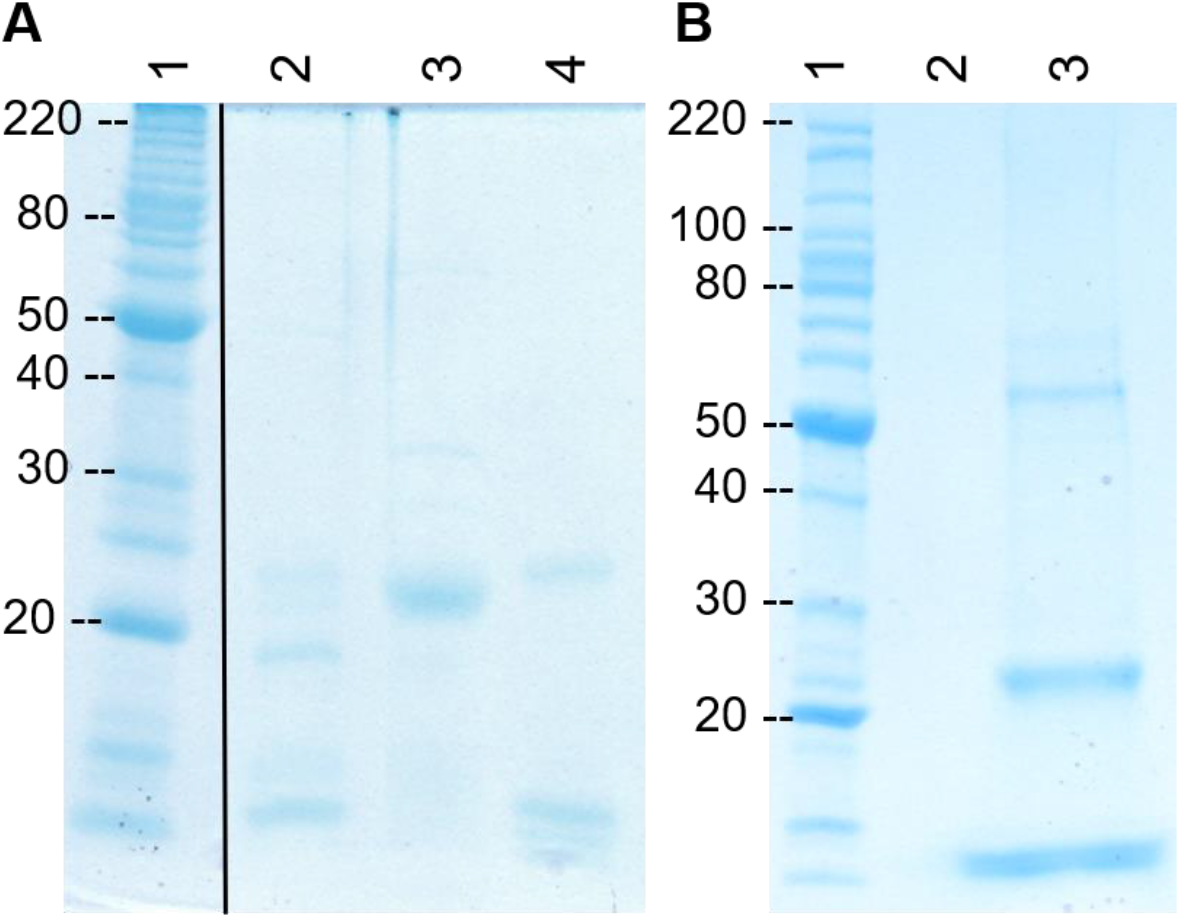
Proteins of filaments purified on cesium sucrose gradients were examined by SDS-PAGE. A, proteins of filaments from sunflower (lane 2), gerbera (lane 3), and zinnia (lane 4). B, proteins from a different filament preparation than shown in A from gerbera (lane 3). No sample was loaded in lane 2.

N-terminal protein sequencing by Edman degradation was performed on the proteins in filament preparations from chrysanthemum, gerbera, and zinnia (Figure 3). The peptide sequences obtained were identical within a species and nearly-identical between the species (Table 2). A tBLASTn search of the EST sequences from chrysanthemum (taxid: 13422) in GenBank revealed that the N-terminal sequences had 100% identity (chrysanthemum 13/13 AA, zinnia 12/12 AA), or nearly perfect identity (gerbera, 88% 15/17 AA and 93% positive, 16/17 AA), with predicted N-terminus of mature kunitz-type protease inhibitors (KTI) from chrysanthemum (accession numbers DK939406.1) suggesting that filaments are composed of mature KTI.

**Figure 3.**
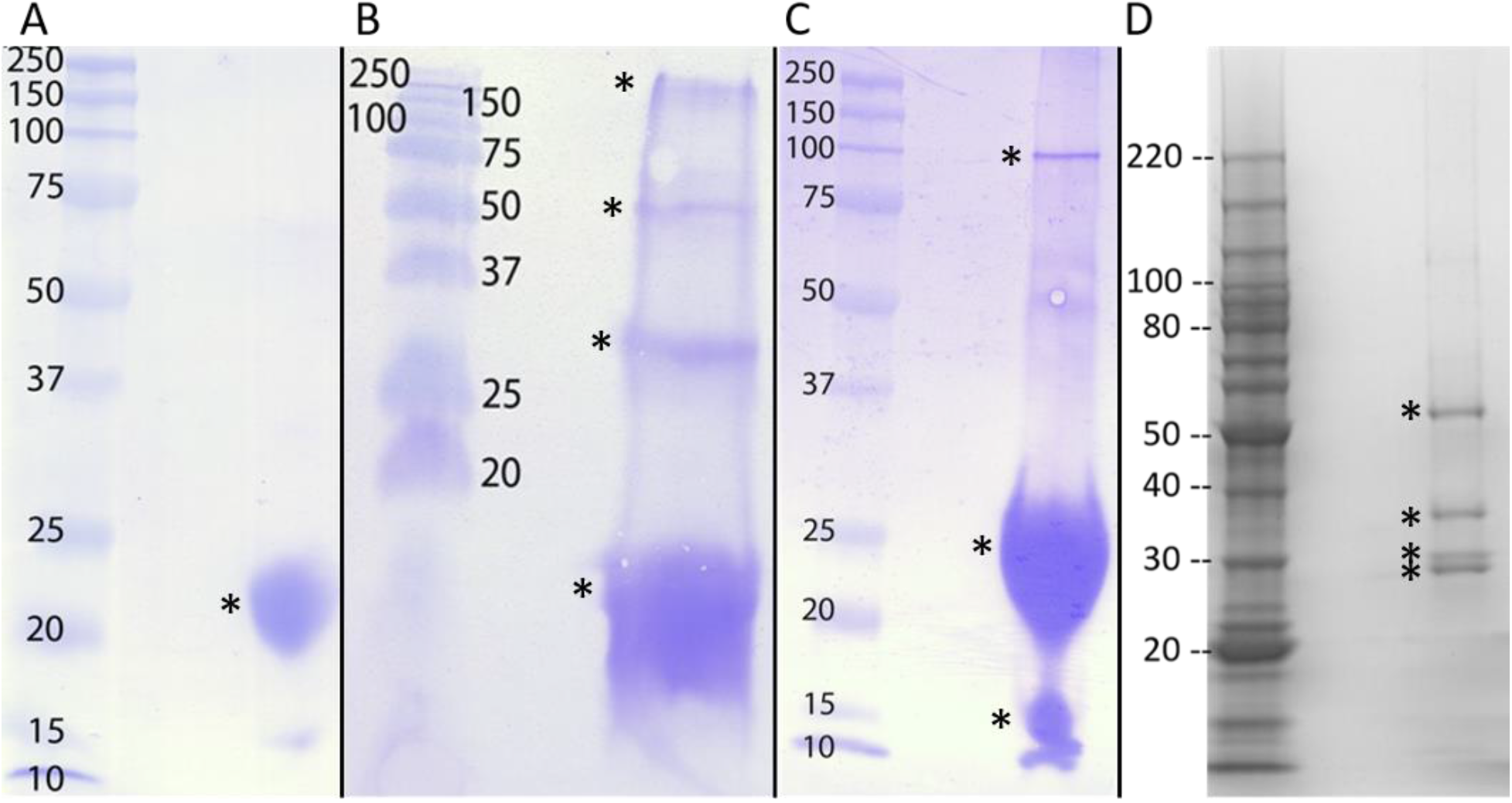
SDS-PAGE of proteins in filament preparations that were sequenced by Edman degradation (A-C) or CID MS/MS (D). The preparations are from gerbera (A), zinnia (B), chrysanthemum (C) or sunflower (D). The bands marked with a * were sequenced.

**Table 2.** N-terminal protein sequencing results for the proteins observed by SDS-PAGE from purified filaments.

|  |  |  | N-Terminal protein sequencing results |  |  |
| --- | --- | --- | --- | --- | --- |
| Band size (kDa) |  | Plant | <u>chrysanthemum</u> | <u>gerbera</u> | <u>zinnia</u> |
|  | 12 | Major | ADVIYD (S+S ' ) AGNKLL |  | ADVIYD (S+S ' ) AGNKL |
|  |  | Minor | GFK (S+S ' ) ATIGNE (S+S ' ) NR |  | VIYD (S+S ' ) K (G/V) (N/V) KLLR |
|  | 22 | Major | ADVIYD (S+S ' ) AGNKLL | (S+S ' ) DVIYD (S+S ' ) DGNKLL (S+S ' ) GVP | ADVIYD (S+S ' ) AGNKL |
|  |  | Minor |  | (S+S ' ) DVIYD (S+S ' ) DGNKLL (S+S ' ) (R/T) VNT |  |
|  | 55 | Major |  |  | ADVIYDSAGNKL |
|  |  | Minor |  |  | SDVIYDSAGNKL |
|  | 100 | Major |  |  | ADVIYDSAGNKL |
|  |  | Minor |  |  | VIYDSKGNKLLL |

To test this hypothesis, we decided to sequence sunflower filament proteins using collision-induced dissociation tandem mass spectrometry (CID MS/MS). At the time we did this the sunflower genome was not yet available and there was no complete cDNA sequence for the KTI most closely related to the chrysanthemum sequence. Therefore, we cloned and sequenced the complete coding region of the related sunflower KTI. The 678 bp product was deposited in GenBank as accession number KY039997.01. The translated amino acid sequence was classified as an STI domain-containing protein (domain architecture ID 11087211) with specific hits to Kunitz_legume (pfam00197) and the STI superfamily (cl11466) proteins. The AA sequence was a perfect match for the chrysanthemum and zinnia N-terminal sequence and 94% match to gerbera.

Sunflower filament proteins were resolved by SDS-PAGE and the four proteins marked in Figure 3D were cut out of the gel, digested with trypsin, and sequences of the resulting peptides were obtained by CID MS/MS analysis and compared to the sunflower KTI sequence (Figures 4 and 5, and supplementary Figures 1-4). The top sequence hit from each of the trypsin fragments had 100% identity with predicted fragments from the cloned sunflower KTI supporting that the sunflower filaments are composed of KTI. None of the sequences mapped to the signal sequence supporting the hypothesis that filaments are composed of the mature protein.

**Figure 4.**
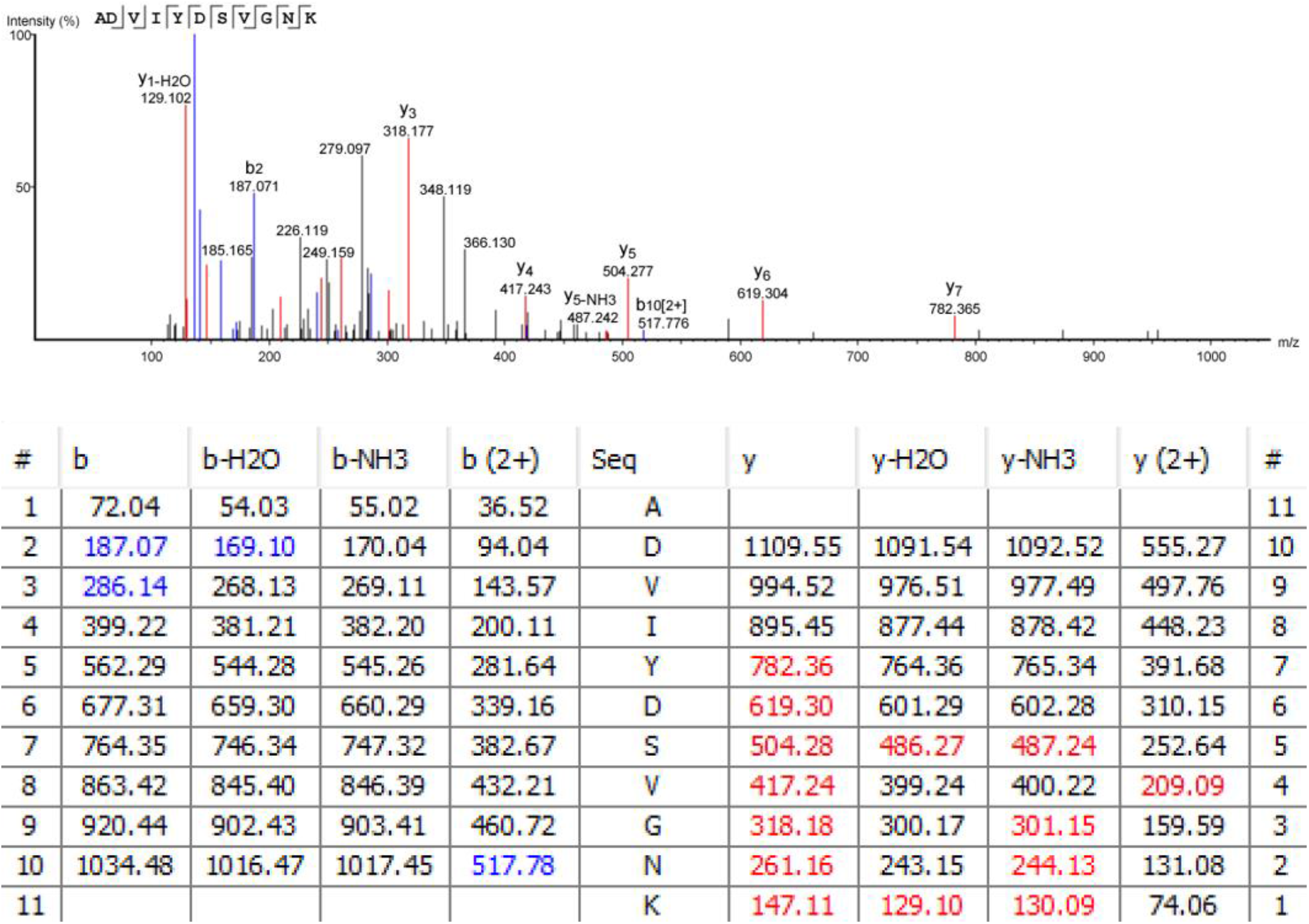
Tandem mass spectra and ion series for the peptide ADVIYDSVGNK from the smallest sunflower protein of ≈28 kDa. The table show the predicted ions. The b and y ions present in the spectrum are in blue and red, respectively.

**Figure 5.**
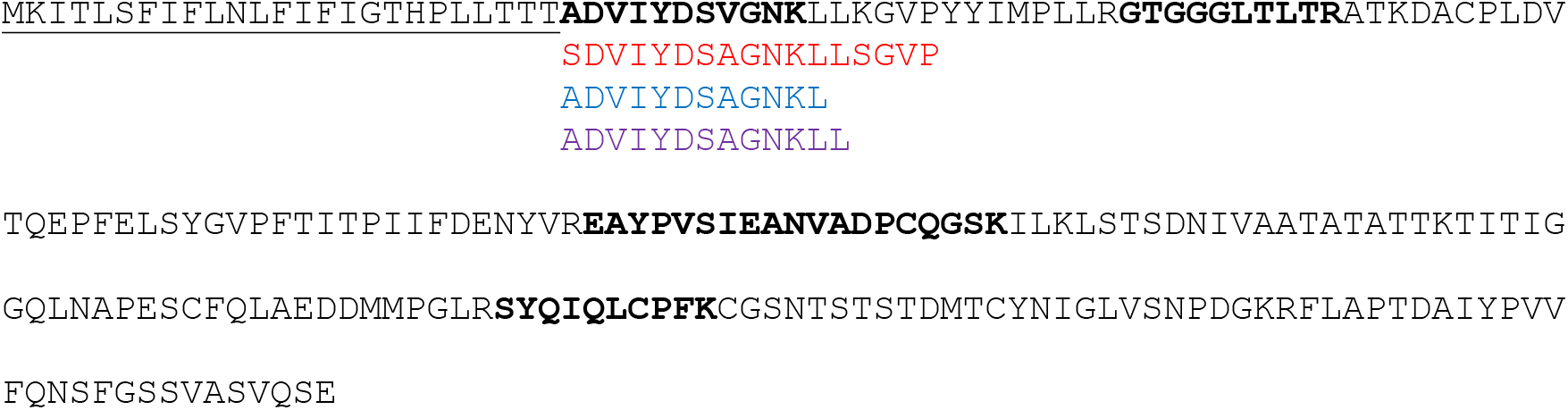
The N-terminal sequences of gerbera (red, top), zinnia, (blue, middle) and chrysanthemum (purple, bottom) filament protein aligned with the sunflower KTI sequence (KY039997) (black). The predicted signal sequence is underlined and the peptides identified by CID MS/MS are in bold. Note that consistent with the protein being the mature sunflower KTI, the first sunflower sequence begins at the predicted signal sequence cleave site rather than a trypsin site.

Several KTIs are glycoproteins (Ee et al. 2009). Therefore, we used glycostaining to determine if gerbera filament proteins were glycosylated (Figure 6). This staining showed that the major 22 kDa protein and a smaller ≈10 kDa protein were glycosylated.

**Figure 6.**
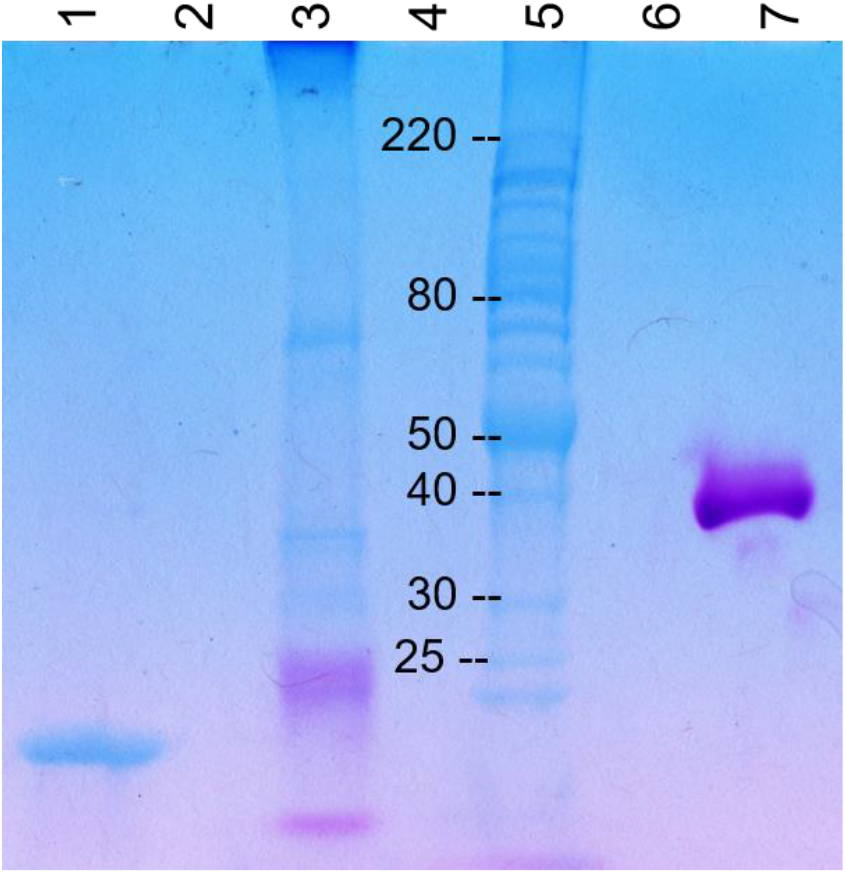
The major gerbera filament protein is glycosylated. Filament proteins were subjected to SDS PAGE and stained to detect glycoproteins. Lane 1 contains purified tobacco mosaic virus as a negative control, lane 3 purified gerbera filaments, lane 5 Benchmark^TM^ protein ladder, lane 7 horseradish peroxidase as a positive control provided with the kit. No samples were loaded into lanes 2, 4, 6.

## Discussion

In the course of searching for plant viruses in members of the *Asteraceae* filamentous structures were identified. Filaments extracted from chrysanthemum (*Chrysanthemum* spp.), coneflower (*Echinacea* spp.), gerbera (*Gerbera jamesonii, G. hybrida*), sunflower (*Helianthus annuus*), and zinnia (*Zinnia hybrida*) had mean diameters ranging from 7.6 to 8.9 nm and were of variable length ranging up to several thousand nm. These filaments are not similar to virions of known plant viruses and we were not able to identify any associated virus nucleic acid. Therefore, we decided to characterize the filament proteins.

When subjected to SDS-PAGE the filament preparations from chrysanthemum, gerbera, sunflower and zinnia contained several proteins. The initial N-terminal protein sequencing of major filament associated proteins from three species (gerbera, zinnia, chrysanthemum) all yielded nearly the same sequence, which matched a predicted mature KTI sequence from chrysanthemum. The nearly identical N-terminal sequence from three different species indicates the filamentous forming protein may be conserved across *Asteraceae*. The predicted size of the mature sunflower KTI is 21.5kDa, which corresponds well to the size of the most abundant protein in each filament preparation. Based on these results we believe that the filaments are composed of the mature KTI and that the other proteins present in the filaments are KTI breakdown products or KTI aggregates. This hypothesis is supported by the fact that the abundance of many of these proteins relative to the mature KTI varied between filament preparations. This effect may be observed with the preparations from zinnia which in one experiment yielded proteins of <10 and 22 kDa while another experiment yielded proteins of <10, 22, and 55 kDa (Fig 1 A-lane 3 and B-lane 3). In addition, the sequence of proteins larger than mature KTI from zinnia (Table 2) and sunflower (supplemental Figures 1-4) indicated they were KTI.

We cloned and sequenced a sunflower gene with high similarity to chrysanthemum KTI and compared the predicted protein sequence to the major filament proteins of sunflower fibril as determined by CID MS/MS. The sequence of four of the trypsin fragments, that were present in each of the protein bands sequenced, had 100% identity with the translated, sunflower KTI and covers 25% of the total sequence (51 out of 200 amino acids). The presence of KTI filaments in the four species examined suggest that these filaments may occur in all or many of the *Asteraceae*.

Several KTIs are glycoproteins (Ee et al. 2009, Paiva et al. 2003, Macedo et al. 2003, Azarkan et al. 2006, do Soccoro et al. 2002) and KTI in filaments was glycosylated. This suggests that filaments form from the mature KTI after it has exited the secretory pathway, perhaps in the protein bodies.

Although not widely studied, proteins and enzymes from animals, fungi and plants are known to assemble into filaments (Park & Horton 2019). Assembly into filaments can change the enzymatic properties (Park & Horton 2019). In plants β-glucosidase, ribonucleotide reductase, and IRE1 form filaments (Park & Horton 2019). This raises the question does KTI exist both in both filamentous and non-filamentous forms and are the properties of the two forms different? An alternative is that forming filaments is a way to pack KTI in protein bodies. Since KTI are involved in defense against herbivory it is also possible that filament formation is important to the activity or stability of the filaments in the herbivore.

Another question that should be addressed is if the filaments have a function in plants. We identified filaments in plants showing “disease symptoms” including flower doubling and leaf curling and distortion and have noted that the abundance of filaments was highly variable. Plants without visible symptoms can contain very low, and sometimes undetectable levels of filaments. Interestingly, some KTIs have demonstrated roles in leaf shape, shoot growth and root growth (Islam et al 2015; Li et al 2008). Further studies are needed to determine if filament abundance is correlated with and can perhaps cause flower doubling, leaf curling, and distortion.

### Accession Numbers

Sequence data from this article can be found in GenBank under accession number

### Supplementary Data files

Supplementary Figures

## Acknowledgements

HATCH/MAES projects MIN -071-019 and MIN-71-041, Fred C. Gloeckner Foundation Inc., MnDRIVE Global Food Initiative, University of Minnesota Doctoral Dissertation Fellowship,

## Conflict of Interest Statement

The authors have no conflicts of interest to declare.

## Author Contributions

SAB- designed the research; performed research; analyzed data; wrote the paper

NEO- designed the research; performed research; analyzed data; wrote the paper

BEL- designed the research; performed research; analyzed data; edited the paper

## Notes

### Competing Interest Statement

The authors have declared no competing interest.

