## Supplementary Figures for "Identification of a filamentous form of kunitz protease inhibitor in *Asteraceae*"

Supplemental Figure 1. Tandem mass spectra and ion series for all peptides obtained from the ≈28 kDa sunflower filament protein. The tables show the predicted ions. The b and y ions present in the spectrum are in blue and red, respectively. Peptides are ordered from N to C-terminus.

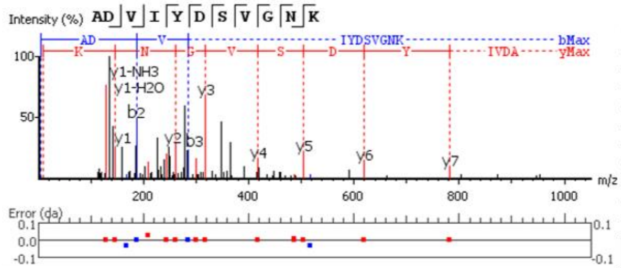


| # | b | b-H2O | b-NH3 | b (2+) | Seq | y | y-H2O | y-NH3 | y (2+) | # |
|---|---|---|---|---|---|---|---|---|---|---|
| 1 | 72.04 | 54.03 | 55.02 | 36.52 | A | | | | | 11 |
| 2 | 187.07 | 169.10 | 170.04 | 94.04 | D | 1109.55 | 1091.54 | 1092.52 | 555.27 | 10 |
| 3 | 286.14 | 268.13 | 269.11 | 143.57 | V | 994.52 | 976.51 | 977.49 | 497.76 | 9 |
| 4 | 399.22 | 381.21 | 382.20 | 200.11 | I | 895.45 | 877.44 | 878.42 | 448.23 | 8 |
| 5 | 562.29 | 544.28 | 545.26 | 281.64 | Y | 782.36 | 764.36 | 765.34 | 391.68 | 7 |
| 6 | 677.31 | 659.30 | 660.29 | 339.16 | D | 619.30 | 601.29 | 602.28 | 310.15 | 6 |
| 7 | 764.35 | 746.34 | 747.32 | 382.67 | S | 504.28 | 486.27 | 487.24 | 252.64 | 5 |
| 8 | 863.42 | 845.40 | 846.39 | 432.21 | V | 417.24 | 399.24 | 400.22 | 209.09 | 4 |
| 9 | 920.44 | 902.43 | 903.41 | 460.72 | G | 318.18 | 300.17 | 301.15 | 159.59 | 3 |
| 10 | 1034.48 | 1016.47 | 1017.45 | 517.78 | N | 261.16 | 243.15 | 244.13 | 131.08 | 2 |
| 11 | | | | | K | 147.11 | 129.10 | 130.09 | 74.06 | 1 |

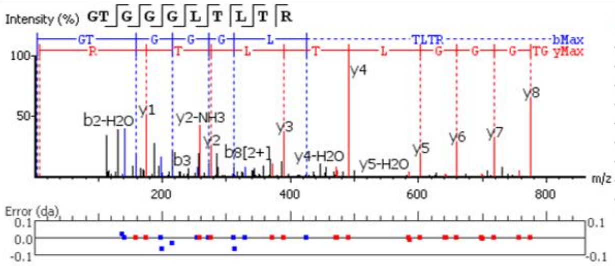


| # | b | b-H2O | b-NH3 | b (2+) | Seq | y | y-H2O | y-NH3 | y (2+) | # |
|---|---|---|---|---|---|---|---|---|---|---|
| 1 | 58.03 | 40.02 | 41.00 | 29.51 | G | | | | | 10 |
| 2 | 159.08 | 141.07 | 142.05 | 80.04 | T | 875.49 | 857.48 | 858.47 | 438.25 | 9 |
| 3 | 216.13 | 198.09 | 199.14 | 108.55 | G | 774.45 | 756.44 | 757.42 | 387.72 | 8 |
| 4 | 273.12 | 255.11 | 256.09 | 137.03 | G | 717.42 | 699.42 | 700.41 | 359.21 | 7 |
| 5 | 330.14 | 312.13 | 313.19 | 165.57 | G | 660.40 | 642.40 | 643.38 | 330.70 | 6 |
| 6 | 443.23 | 425.22 | 426.20 | 222.11 | L | 603.38 | 585.37 | 586.37 | 302.19 | 5 |
| 7 | 544.27 | 526.26 | 527.25 | 272.64 | T | 490.30 | 472.29 | 473.27 | 245.65 | 4 |
| 8 | 657.36 | 639.35 | 640.33 | 329.18 | L | 389.25 | 371.24 | 372.22 | 195.13 | 3 |
| 9 | 758.40 | 740.39 | 741.38 | 379.70 | T | 276.17 | 258.16 | 259.14 | 138.58 | 2 |
| 10 | | | | | R | 175.12 | 157.11 | 158.09 | 88.06 | 1 |

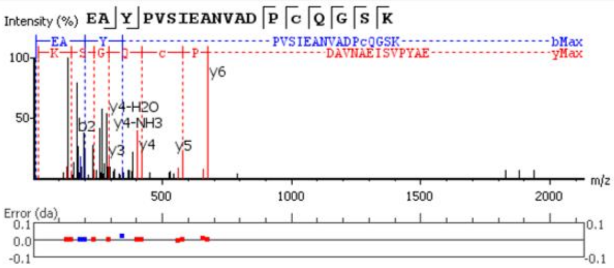


| # | b | b-H2O | b-NH3 | b (2+) | Seq | y | y-H2O | y-NH3 | y (2+) | # |
|---|---|---|---|---|---|---|---|---|---|---|
| 1 | 130.05 | 112.04 | 113.02 | 65.53 | E | | | | | 19 |
| 2 | 201.09 | 183.08 | 184.06 | 101.04 | A | 1905.90 | 1887.89 | 1888.87 | 953.45 | 18 |
| 3 | 364.15 | 346.12 | 347.12 | 182.58 | Y | 1834.86 | 1816.85 | 1817.84 | 917.93 | 17 |
| 4 | 461.20 | 443.19 | 444.18 | 231.10 | P | 1671.80 | 1653.79 | 1654.77 | 836.40 | 16 |
| 5 | 560.27 | 542.26 | 543.24 | 280.64 | V | 1574.75 | 1556.74 | 1557.72 | 787.87 | 15 |
| 6 | 647.30 | 629.29 | 630.28 | 324.15 | S | 1475.68 | 1457.67 | 1458.65 | 738.34 | 14 |
| 7 | 760.39 | 742.38 | 743.36 | 380.69 | I | 1388.65 | 1370.64 | 1371.62 | 694.82 | 13 |
| 8 | 889.43 | 871.42 | 872.40 | 445.22 | E | 1275.56 | 1257.55 | 1258.54 | 638.28 | 12 |
| 9 | 960.47 | 942.46 | 943.44 | 480.73 | A | 1146.52 | 1128.51 | 1129.49 | 573.76 | 11 |
| 10 | 1074.51 | 1056.50 | 1057.48 | 537.76 | N | 1075.48 | 1057.47 | 1058.46 | 538.24 | 10 |
| 11 | 1173.58 | 1155.57 | 1156.55 | 587.29 | V | 961.44 | 943.43 | 944.41 | 481.22 | 9 |
| 12 | 1244.62 | 1226.61 | 1227.59 | 622.81 | A | 862.37 | 844.36 | 845.35 | 431.69 | 8 |
| 13 | 1359.64 | 1341.63 | 1342.62 | 680.32 | D | 791.34 | 773.32 | 774.31 | 396.17 | 7 |
| 14 | 1456.70 | 1438.69 | 1439.67 | 728.85 | P | 676.31 | 658.29 | 659.27 | 338.65 | 6 |
| 15 | 1616.73 | 1598.72 | 1599.70 | 808.86 | C(+57.02) | 579.25 | 561.24 | 562.24 | 290.13 | 5 |
| 16 | 1744.79 | 1726.77 | 1727.76 | 872.89 | Q | 419.22 | 401.21 | 402.20 | 210.11 | 4 |
| 17 | 1801.81 | 1783.80 | 1784.78 | 901.40 | G | 291.17 | 273.16 | 274.14 | 146.08 | 3 |
| 18 | 1888.84 | 1870.83 | 1871.81 | 944.92 | S | 234.14 | 216.13 | 217.12 | 117.57 | 2 |
| 19 | | | | | K | 147.11 | 129.10 | 130.09 | 74.06 | 1 |

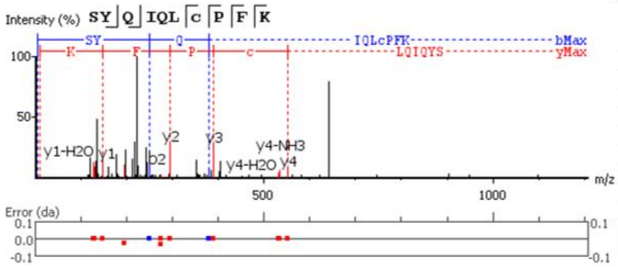


| # | b | b-H2O | b-NH3 | b (2+) | Seq | y | y-H2O | y-NH3 | y (2+) | # |
|---|---|---|---|---|---|---|---|---|---|---|
| 1 | 88.04 | 70.03 | 71.01 | 44.52 | S | | | | | 10 |
| 2 | 251.10 | 233.09 | 234.08 | 126.05 | Y | 1196.61 | 1178.60 | 1179.59 | 598.81 | 9 |
| 3 | 379.16 | 361.15 | 362.13 | 190.08 | Q | 1033.55 | 1015.54 | 1016.52 | 517.27 | 8 |
| 4 | 492.25 | 474.24 | 475.22 | 246.62 | I | 905.49 | 887.48 | 888.46 | 453.25 | 7 |
| 5 | 620.30 | 602.29 | 603.28 | 310.65 | Q | 792.41 | 774.40 | 775.38 | 396.70 | 6 |
| 6 | 733.39 | 715.38 | 716.36 | 367.19 | L | 664.35 | 646.34 | 647.32 | 332.67 | 5 |
| 7 | 893.42 | 875.41 | 876.39 | 447.21 | C(+57.02) | 551.26 | 533.25 | 534.24 | 276.17 | 4 |
| 8 | 990.47 | 972.46 | 973.44 | 495.74 | P | 391.23 | 373.22 | 374.21 | 196.14 | 3 |
| 9 | 1137.54 | 1119.53 | 1120.51 | 569.27 | F | 294.18 | 276.17 | 277.15 | 147.59 | 2 |
| 10 | | | | | K | 147.11 | 129.10 | 130.09 | 74.06 | 1 |

Supplemental Data. Bratsch et al. 1-22-2021. “Identification of a filamentous form of kunitz protease inhibitor in *Asteraceae*”

Supplemental Figure 2. Tandem mass spectra and ion series for all peptides obtained from the ≈30 kDa sunflower filament protein. The tables show the predicted ions. The b and y ions present in the spectrum are in blue and red, respectively. Peptides are ordered from N to C-terminus.

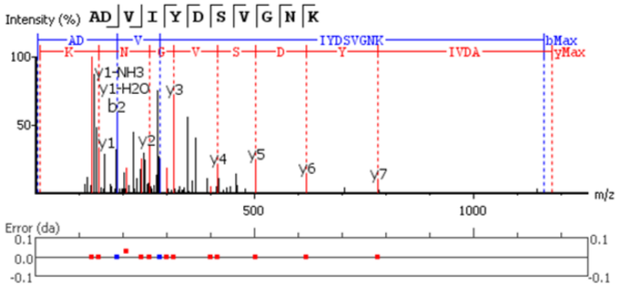


| # | b | b-H2O | b-NH3 | b (2+) | Seq | y | y-H2O | y-NH3 | y (2+) | # |
|---|---|---|---|---|---|---|---|---|---|---|
| 1 | 72.04 | 54.03 | 55.02 | 36.52 | A | | | | | 11 |
| 2 | 187.07 | 169.06 | 170.04 | 94.04 | D | 1109.55 | 1091.54 | 1092.52 | 555.27 | 10 |
| 3 | 286.14 | 268.13 | 269.11 | 143.57 | V | 994.52 | 976.51 | 977.49 | 497.76 | 9 |
| 4 | 399.22 | 381.21 | 382.20 | 200.11 | I | 895.45 | 877.44 | 878.42 | 448.23 | 8 |
| 5 | 562.29 | 544.28 | 545.26 | 281.64 | Y | 782.37 | 764.36 | 765.34 | 391.68 | 7 |
| 6 | 677.31 | 659.30 | 660.29 | 339.16 | D | 619.30 | 601.29 | 602.28 | 310.15 | 6 |
| 7 | 764.35 | 746.34 | 747.32 | 382.67 | S | 504.28 | 486.27 | 487.25 | 252.64 | 5 |
| 8 | 863.42 | 845.40 | 846.39 | 432.21 | V | 417.25 | 399.24 | 400.22 | 209.09 | 4 |
| 9 | 920.44 | 902.43 | 903.41 | 460.72 | G | 318.18 | 300.17 | 301.15 | 159.59 | 3 |
| 10 | 1034.48 | 1016.47 | 1017.45 | 517.74 | N | 261.16 | 243.15 | 244.13 | 131.08 | 2 |
| 11 | | | | | K | 147.11 | 129.10 | 130.09 | 74.06 | 1 |

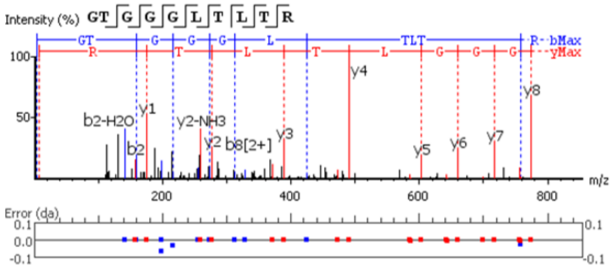


| # | b | b-H2O | b-NH3 | b (2+) | Seq | y | y-H2O | y-NH3 | y (2+) | # |
|---|---|---|---|---|---|---|---|---|---|---|
| 1 | 58.03 | 40.02 | 41.00 | 29.51 | G | | | | | 10 |
| 2 | 159.08 | 141.07 | 142.05 | 80.04 | T | 875.49 | 857.48 | 858.47 | 438.25 | 9 |
| 3 | 216.13 | 198.09 | 199.14 | 108.55 | G | 774.45 | 756.43 | 757.43 | 387.72 | 8 |
| 4 | 273.12 | 255.11 | 256.09 | 137.06 | G | 717.43 | 699.41 | 700.40 | 359.21 | 7 |
| 5 | 330.14 | 312.13 | 313.11 | 165.57 | G | 660.40 | 642.39 | 643.39 | 330.70 | 6 |
| 6 | 443.23 | 425.22 | 426.20 | 222.11 | L | 603.38 | 585.37 | 586.37 | 302.19 | 5 |
| 7 | 544.27 | 526.26 | 527.25 | 272.64 | T | 490.30 | 472.29 | 473.27 | 245.65 | 4 |
| 8 | 657.36 | 639.35 | 640.33 | 329.18 | L | 389.25 | 371.24 | 372.22 | 195.13 | 3 |
| 9 | 758.44 | 740.39 | 741.38 | 379.70 | T | 276.17 | 258.16 | 259.14 | 138.58 | 2 |
| 10 | | | | | R | 175.12 | 157.11 | 158.09 | 88.06 | 1 |

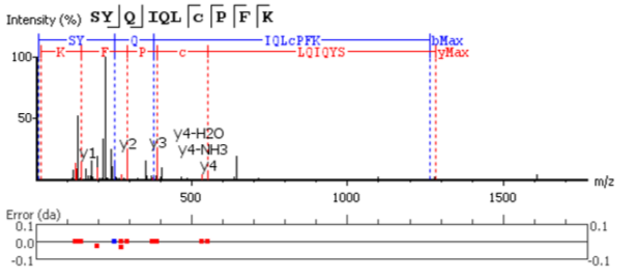


| # | b | b-H2O | b-NH3 | b (2+) | Seq | y | y-H2O | y-NH3 | y (2+) | # |
|---|---|---|---|---|---|---|---|---|---|---|
| 1 | 88.04 | 70.03 | 71.01 | 44.52 | S | | | | | 10 |
| 2 | 251.10 | 233.09 | 234.08 | 126.05 | Y | 1196.61 | 1178.60 | 1179.59 | 598.81 | 9 |
| 3 | 379.16 | 361.15 | 362.13 | 190.08 | Q | 1033.55 | 1015.54 | 1016.52 | 517.27 | 8 |
| 4 | 492.25 | 474.24 | 475.22 | 246.62 | I | 905.49 | 887.48 | 888.46 | 453.25 | 7 |
| 5 | 620.30 | 602.29 | 603.28 | 310.65 | Q | 792.41 | 774.40 | 775.38 | 396.70 | 6 |
| 6 | 733.39 | 715.38 | 716.36 | 367.19 | L | 664.35 | 646.34 | 647.32 | 332.67 | 5 |
| 7 | 893.42 | 875.41 | 876.39 | 447.21 | C(+57.02) | 551.27 | 533.26 | 534.24 | 276.17 | 4 |
| 8 | 990.47 | 972.46 | 973.44 | 495.74 | P | 391.23 | 373.22 | 374.21 | 196.14 | 3 |
| 9 | 1137.54 | 1119.53 | 1120.51 | 569.27 | F | 294.18 | 276.17 | 277.15 | 147.59 | 2 |
| 10 | | | | | K | 147.11 | 129.10 | 130.09 | 74.06 | 1 |

Supplemental Data. Bratsch et al. 1-22-2021. “Identification of a filamentous form of kunitz protease inhibitor in *Asteraceae*”

Supplemental Figure 3. Tandem mass spectra and ion series for all peptides obtained from the ≈35 kDa sunflower filament protein. The tables show the predicted ions. The b and y ions present in the spectrum are in blue and red, respectively. Peptides are ordered from N to C-terminus.

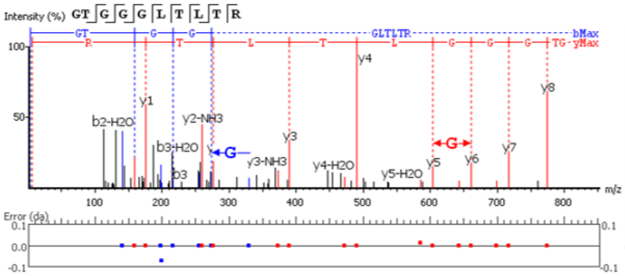


| # | b | b-H2O | b-NH3 | b (2+) | Seq | y | y-H2O | y-NH3 | y (2+) | # |
|---|---|---|---|---|---|---|---|---|---|---|
| 1 | 58.03 | 40.02 | 41.00 | 29.51 | G | | | | | 10 |
| 2 | 159.08 | 141.07 | 142.05 | 80.04 | T | 875.49 | 857.48 | 858.47 | 438.25 | 9 |
| 3 | 216.10 | 198.09 | 199.15 | 108.55 | G | 774.45 | 756.44 | 757.42 | 387.72 | 8 |
| 4 | 273.12 | 255.11 | 256.09 | 137.06 | G | 717.42 | 699.42 | 700.40 | 359.21 | 7 |
| 5 | 330.14 | 312.13 | 313.11 | 165.57 | G | 660.40 | 642.40 | 643.38 | 330.70 | 6 |
| 6 | 443.23 | 425.21 | 426.20 | 222.11 | L | 603.39 | 585.36 | 586.36 | 302.19 | 5 |
| 7 | 544.27 | 526.26 | 527.25 | 272.64 | T | 490.30 | 472.29 | 473.27 | 245.65 | 4 |
| 8 | 657.36 | 639.35 | 640.33 | 329.18 | L | 389.25 | 371.24 | 372.23 | 195.13 | 3 |
| 9 | 758.40 | 740.39 | 741.38 | 379.70 | T | 276.17 | 258.16 | 259.14 | 138.58 | 2 |
| 10 | | | | | R | 175.12 | 157.11 | 158.09 | 88.06 | 1 |

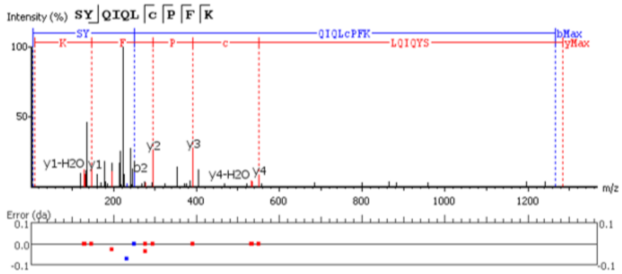


| # | b | b-H2O | b-NH3 | b (2+) | Seq | y | y-H2O | y-NH3 | y (2+) | # |
|---|---|---|---|---|---|---|---|---|---|---|
| 1 | 88.04 | 70.03 | 71.01 | 44.52 | S | | | | | 10 |
| 2 | 251.10 | 233.16 | 234.08 | 126.05 | Y | 1196.61 | 1178.60 | 1179.59 | 598.81 | 9 |
| 3 | 379.16 | 361.15 | 362.13 | 190.08 | Q | 1033.55 | 1015.54 | 1016.52 | 517.27 | 8 |
| 4 | 492.25 | 474.24 | 475.22 | 246.62 | I | 905.49 | 887.48 | 888.46 | 453.25 | 7 |
| 5 | 620.30 | 602.29 | 603.28 | 310.65 | Q | 792.41 | 774.40 | 775.38 | 396.70 | 6 |
| 6 | 733.39 | 715.38 | 716.36 | 367.19 | L | 664.35 | 646.34 | 647.32 | 332.67 | 5 |
| 7 | 893.42 | 875.41 | 876.39 | 447.21 | C(+57.02) | 551.27 | 533.26 | 534.24 | 276.17 | 4 |
| 8 | 990.47 | 972.46 | 973.44 | 495.74 | P | 391.23 | 373.22 | 374.21 | 196.14 | 3 |
| 9 | 1137.54 | 1119.53 | 1120.51 | 569.27 | F | 294.18 | 276.17 | 277.15 | 147.59 | 2 |
| 10 | | | | | K | 147.11 | 129.10 | 130.09 | 74.06 | 1 |

Supplemental Data. Bratsch et al. 1-22-2021. “Identification of a filamentous form of kunitz protease inhibitor in *Asteraceae*”

Supplemental Figure 4. Tandem mass spectra and ion series for all peptides obtained from the ≈55 kDa sunflower filament protein. The tables show the predicted ions. The b and y ions present in the spectrum are in blue and red, respectively. Peptides are ordered from N to C-terminus.

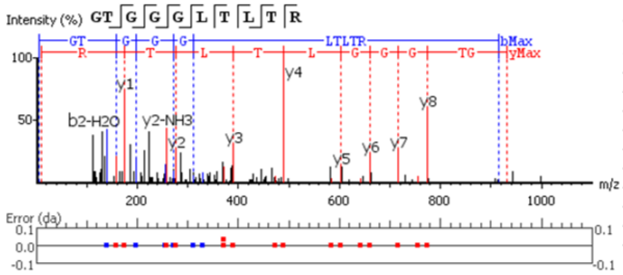


| # | b | b-H2O | b-NH3 | b (2+) | Seq | y | y-H2O | y-NH3 | y (2+) | # |
|---|---|---|---|---|---|---|---|---|---|---|
| 1 | 58.03 | 40.02 | 41.00 | 29.51 | G | | | | | 10 |
| 2 | 159.08 | 141.07 | 142.05 | 80.04 | T | 875.49 | 857.48 | 858.47 | 438.25 | 9 |
| 3 | 216.10 | 198.09 | 199.07 | 108.55 | G | 774.45 | 756.44 | 757.42 | 387.72 | 8 |
| 4 | 273.12 | 255.11 | 256.09 | 137.06 | G | 717.43 | 699.41 | 700.40 | 359.21 | 7 |
| 5 | 330.14 | 312.13 | 313.11 | 165.57 | G | 660.40 | 642.40 | 643.38 | 330.70 | 6 |
| 6 | 443.23 | 425.21 | 426.20 | 222.11 | L | 603.38 | 585.37 | 586.36 | 302.19 | 5 |
| 7 | 544.27 | 526.26 | 527.25 | 272.64 | T | 490.30 | 472.29 | 473.27 | 245.65 | 4 |
| 8 | 657.36 | 639.35 | 640.33 | 329.18 | L | 389.25 | 371.21 | 372.22 | 195.13 | 3 |
| 9 | 758.40 | 740.39 | 741.38 | 379.70 | T | 276.17 | 258.16 | 259.14 | 138.58 | 2 |
| 10 | | | | | R | 175.12 | 157.11 | 158.09 | 88.06 | 1 |

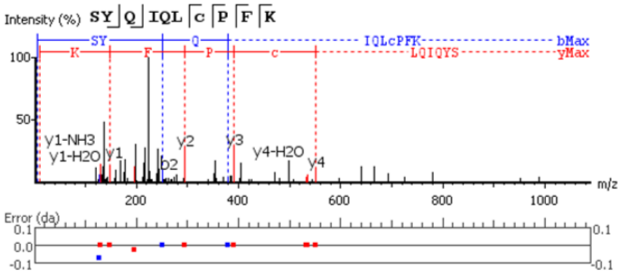


| # | b | b-H2O | b-NH3 | b (2+) | Seq | y | y-H2O | y-NH3 | y (2+) | # |
|---|---|---|---|---|---|---|---|---|---|---|
| 1 | 88.04 | 70.03 | 71.01 | 44.52 | S | | | | | 10 |
| 2 | 251.10 | 233.09 | 234.08 | 126.13 | Y | 1196.61 | 1178.60 | 1179.59 | 598.81 | 9 |
| 3 | 379.16 | 361.15 | 362.13 | 190.08 | Q | 1033.55 | 1015.54 | 1016.52 | 517.27 | 8 |
| 4 | 492.25 | 474.24 | 475.22 | 246.62 | I | 905.49 | 887.48 | 888.46 | 453.25 | 7 |
| 5 | 620.30 | 602.29 | 603.28 | 310.65 | Q | 792.41 | 774.40 | 775.38 | 396.70 | 6 |
| 6 | 733.39 | 715.38 | 716.36 | 367.19 | L | 664.35 | 646.34 | 647.32 | 332.67 | 5 |
| 7 | 893.42 | 875.41 | 876.39 | 447.21 | C(+57.02) | 551.27 | 533.26 | 534.24 | 276.13 | 4 |
| 8 | 990.47 | 972.46 | 973.44 | 495.74 | P | 391.23 | 373.22 | 374.21 | 196.14 | 3 |
| 9 | 1137.54 | 1119.53 | 1120.51 | 569.27 | F | 294.18 | 276.17 | 277.15 | 147.59 | 2 |
| 10 | | | | | K | 147.11 | 129.10 | 130.09 | 74.06 | 1 |
